# TERT-mediated induction of MIR500A contributes to tumor invasiveness by targeting Hedgehog pathway

**DOI:** 10.1101/2020.02.18.954370

**Authors:** Manuel Bernabé-García, Elena Martínez-Balsalobre, Diana García-Moreno, Jesús García-Castillo, Beatriz Revilla-Nuin, Elena Blanco-Alcaina, Victoriano Mulero, Francisca Alcaraz-Pérez, María L. Cayuela

## Abstract

The classical activity of telomerase (TERT) is to maintain telomere homeostasis, ensuring chromosome stability and cellular proliferation. However, increasing evidences of telomere-independent human TERT functions have been lastly obtained. We report here that TERT directly binds to the TCF binding elements (TBE) located upstream the oncomiR *MIR500A* inducing its expression and promoting cancer invasiveness. This function is independent of telomerase activity, since catalytic inactive TERT also induces *MIR500A* expression and telomerase inhibitors directed against TERT, but not to its RNA component *TERC*, inhibit telomerase-induced *MIR500A* expression and cancer invasiveness. Mechanistically, telomerase-induced *MIR500A* down-regulates key genes of the Hedgehog signaling pathway, namely patched 1 (*PTCH1*), Gli family zinc finger 3 (*GLI3*) and cullin 3 (*CUL3*), increasing tumor invasiveness. Our results show a crucial role of the TERT/*MIR500A*/Hedgehog axis is tumor aggressiveness, pointing out to the relevance of inhibiting the extracurricular functions of telomerase to fight cancer.

## Introduction

Human telomerase (TERT) is reactivated in approximately 90% of all cancers while, in approximately 10% of tumor, telomere length is maintained independently of TERT by the homologous recombination (HR)-mediated alternative lengthening of telomeres (ALT) pathway (1). Increasing evidences are revealing non-canonical roles of TERT not only in cancer, but also in several essential cellular functions, via mechanisms independent of telomere maintenance. These novel roles of TERT may provide transformed cells with specific capacities at multiple stages of tumor development (2). Among the numerous non-telomeric biological functions of TERT, it has been demonstrated that TERT acts as a regulatory molecule modulating gene transcription (3-6). However, the non-canonical roles of TERT in cancer and their relevance in its progression and response to therapy remain poorly understood.

miRNAs are endogenous non-coding small RNAs (∼22 nucleotides) that cause post-transcriptional repression or cleavage of target messenger RNAs (mRNAs) by binding to their 3’UTR. Around 50% of all miRNA genes are located within 50 kb in length on the genome and transcribed together as a cluster and frequently shows similar sequence homology in the seed sequence, the region for target recognition, resulting in identical targets for a miRNA cluster (7). Different studies estimated that each miRNA can regulate more than 200 genes (8, 9), implying that miRNAs regulates a large number of biological processes that are frequently altered in many human diseases. Over the past 15 years, a lot of evidence has shown that aberrant miRNAs expression is involved not only in tumorigenesis and metastasis (10) but, in addition, the miRNA expression profile is unique for each cancer type, so blood-based miRNA expression patterns can be used as a non-invasive method for cancer diagnosis (11). In 2014, Drevytska and colleagues showed a positive correlation between the expression of *TERT* and several miRNA (12). Consistent with these results, gastric cancer models revealed that TERT regulates several miRNAs (13). However regulatory mechanisms involved are not fully understood.

Despite significant clinical advancements, the mortality of most solid tumors throughout the world is largely due to the process of metastasis. Metastasis is a highly dynamic process that occurs in multiple steps regulated by several signalling pathways, which remain incompletely understood, especially the initial steps leading to intravasation, when small developing tumors and micrometastasis are not easily detected (14). Thus, there is a crucial need to understand invasive mechanisms and angiogenic programs that facilitate metastasis so that therapeutic strategies can be developed to block disease progression.

Because TERT has a non-canonical role in regulating the expression of genes involved in cancer initiation and dissemination, in this study we sought to identify miRNA regulated by TERT and then used a zebrafish xenograft model (15) to investigate the non-canonical role of TERT in metastasis through the regulation of these miRNAs. We found that TERT directly regulates *MIR500A* by binding the TBE located upstream of this gene, resulting in the inhibition of Hedgehog signaling pathway and increased tumor invasiveness. These results uncover a non-canonical role of TERT in promoting cancer invasiveness and reveal novel targets for therapeutic intervention.

## Results

### Expression of TERT increases tumor cell line invasion

To study the non-canonical functions of telomerase in invasiveness, we stably transfected the cell line SAOS 2, a telomerase-negative osteosarcoma cell line which maintains its telomeres by ALT, with the plasmid pBABE-puro-hTERT. We selected 2 clones with high *TERT* expression (**Fig. S1A**) for zebrafish larvae xenotransplantation assays. Sixty percent of zebrafish larvae injected with the hTERT-SAOS 2 line had cells outside the yolk sac, whereas only 40% of the larvae had invasion after injection of parental cells (pBABE-SAOS 2, **Fig. S1B**). Therefore, TERT expression in SAOS 2 increased their invasiveness.

### TERT regulates the expression of MIR500A

To evaluate whether the TERT-dependent invasiveness of cancer cells is mediated through the regulation of miRNAs, we used a miRNA microarray to analyze the miRNA expression profile in TERT-overexpression conditions and we found that only the oncomiR *MIR500A* was significantly up-regulated (data not shown). We verified this result by RT-qPCR (**Fig. 1A**) and confirmed its correlation with the higher *in vivo* invasiveness of the tumor cells expressing *TERT* (**Fig. 1B**). To confirm if the higher invasiveness of TERT-expressing tumor cells was mediated by *MIR500A*, we manipulated *MIR500A* expression levels in pBABE-SAOS 2 cells by transfecting them with the *pre-MIR500A* or with a PNA-labeled anti-*MIR500A* probe. Strikingly, *MIR500A* overexpression increased the *in vivo* invasive capacity of both control and TERT-expressing SAOS 2 cells (**Figs. 1C, 1D**), while *MIR500A* inhibition specifically reduced the increased invasiveness of hTERT-SAOS 2 cells (**Figs. 1E, 1F**). Similarly, genetic inhibition of TERT in telomerase positive HeLa cells resulted in reduced expression levels of *MIR500A* and impaired invasiveness (**Fig. S2**).

**Figure 1:**
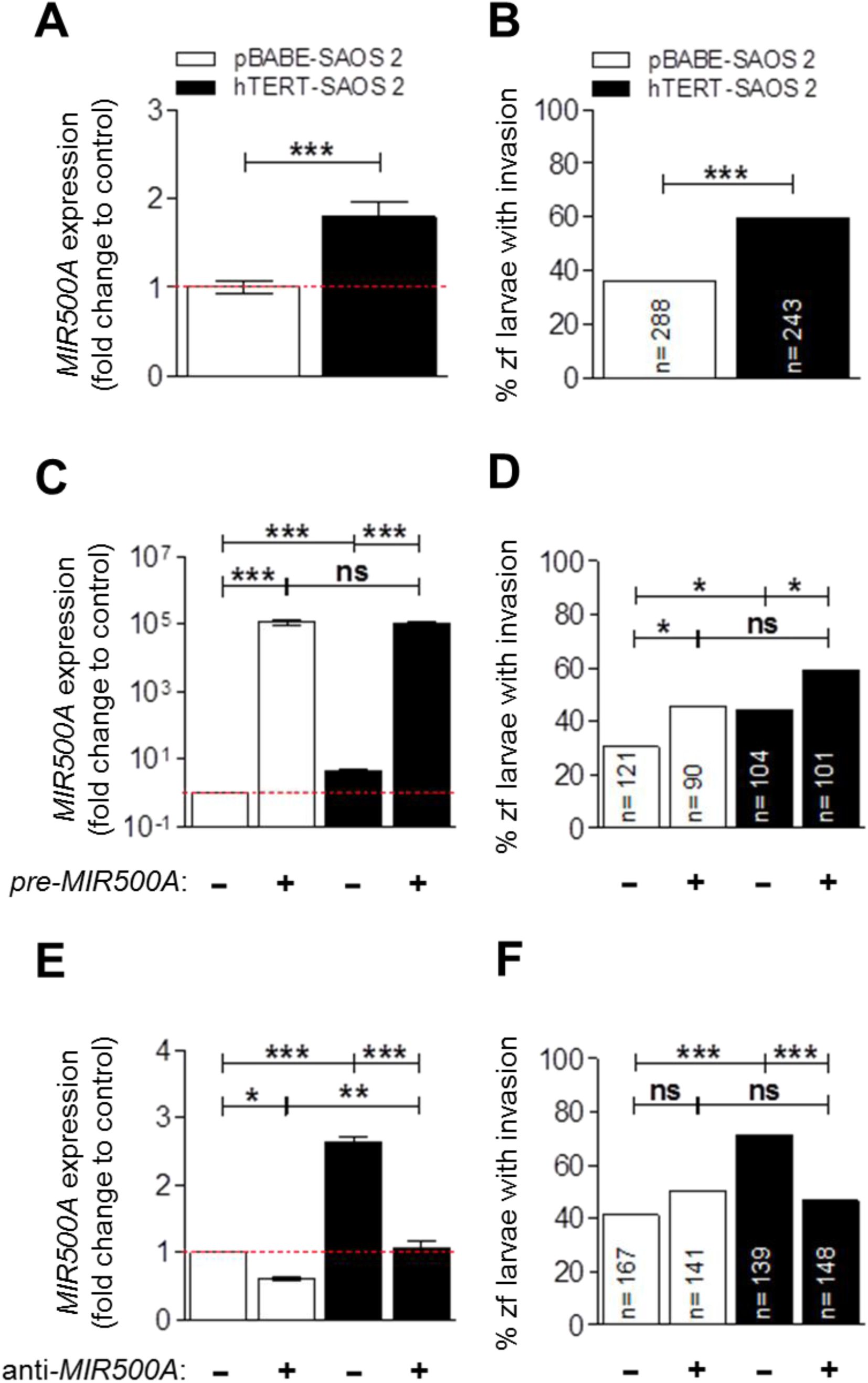
TERT up-regulates the expression of *MIR500A*, which leads to an increase in the *in vivo* invasive capacity. We confirmed the array result determining the *MIR500A* levels in TERT-overexpression conditions by real-time RT-qPCR (**A**). The increased level of *MIR500A* corresponded with an increased *in vivo* invasive capacity of SAOS 2 cells (**B**). Then, we overexpressed and inhibited the *MIR500A* by transient transfection with the *pre-MIR500A* (**C, D**) or with a PNA probe anti-*MIR500A* probe (**E, F**), respectively, in both pBABE- and hTERT-SAOS 2 cells, and we determined the *MIR500A* levels (**C, E**) and the effect on the *in vivo* invasive capacity (**D, F**). In (A, C, E), each bar represents the mean ± SEM from triplicate samples. In (B, D, F), histogram represents the accumulative value of invasion percentage from a total larvae stated in the figure for each treatment. Graphs are representative of three (N= 3) (A, C, E) or the accumulative value of six (N= 6) (B) or four (N= 4) (D, F) different experiments. ns, not significant; *p<0.05; **p<0.01; ***p<0.001 according to Student’s *t-*test (A), ANOVA followed by Tukey’s multiple range test (C, E) and Fisher’s exact test (B, D, F).

### TERT regulates the MIR500 cluster by directly binding to its promoter region

According to the *Ensembl* database (*https://www.ensembl.org*), the oncomiR *MIR500A* is located in a cluster of 8 miRNAs, which is called the MIR500 cluster, into the short arm of the human X chromosome (Xp11.23) and within the intron 3 of the *CLCN5* gene (**Fig. 2A**). Although the majority of intronic miRNAs are transcribed from the same promoter as the host gene, approximately one-third of them are transcribed from independent promoters, enabling separate control of their transcription (16). So we next studied whether the expression of the *CLCN5* gene was affected by TERT and found that *CLCN5* expression is similar in parental and TERT-expressing SAOS 2 cells (**Fig. 2B**). To address the mechanism by which TERT regulates the transcription of *MIR500A*, we cloned into a luciferase reporter plasmid the 2 Kb fragment upstream *MIR500A*, which contains several TCF binding elements (TBE), according to the database PROmiRNA (https://tools4mirs.org/software/other_tools/promirna/). The luciferase reporter assay showed that TERT was able to increase the expression of the reporter driven by the 2 Kb fragment upstream *MIR500A* (**Fig. 2C**). These results were further confirmed in HEK 293 cells, a telomerase positive cell line, where overexpression of hTERT increased while inhibition by siRNA decreased *MIR500A* promoter activity (**Fig. 2D**).

**Figure 2:**
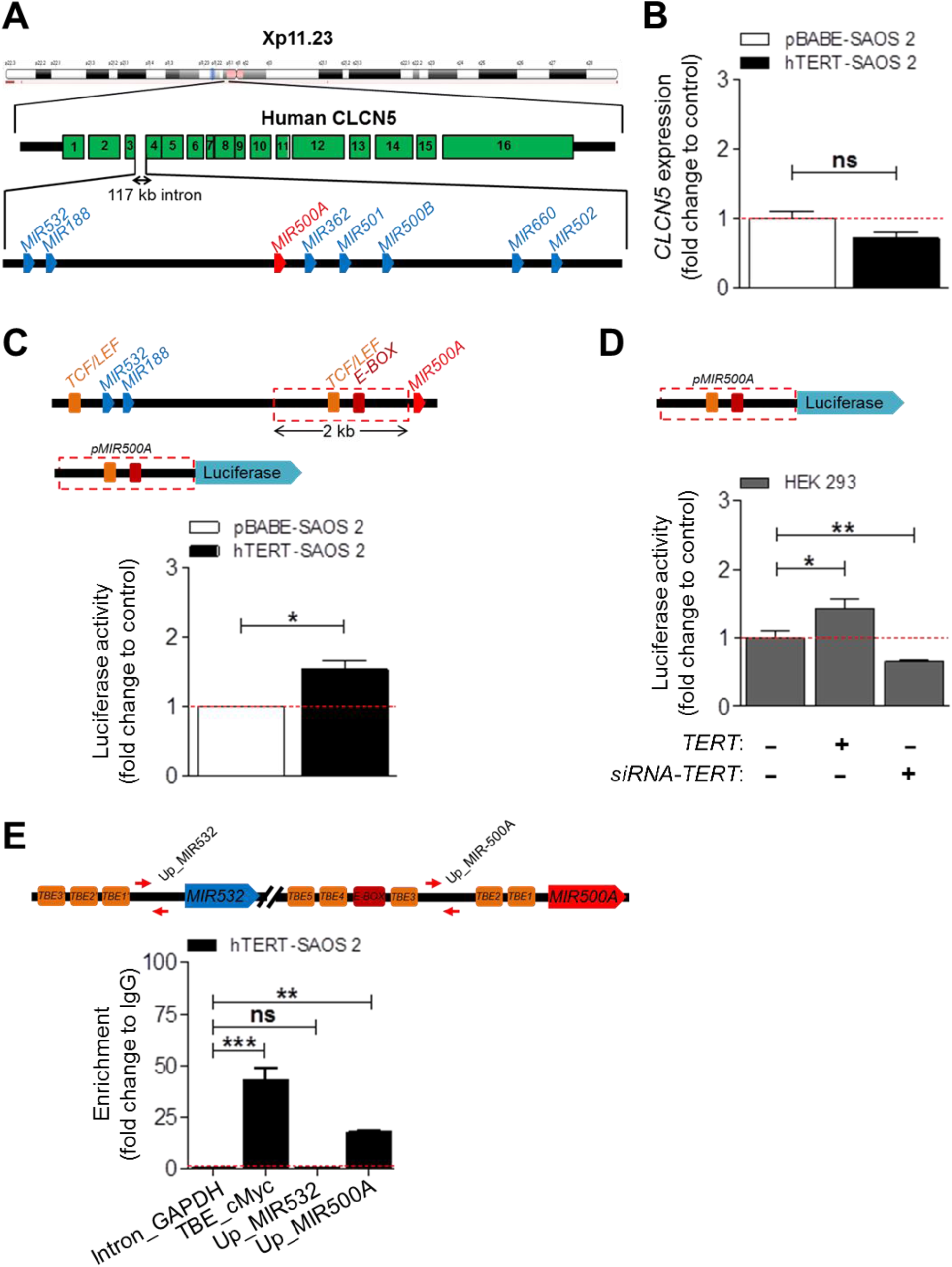
TERT regulates *MIR500A* by direct binding to its promoter region. Schematic representation of the MIR500 cluster according to the *Ensembl* database (**A**). Names are shorter to simplify. We determined the *CLCN5* mRNA levels in TERT-overexpression conditions by real-time RT-qPCR (**B**). Next, we cloned a 2 Kb region upstream the *MIR500A* gene driving the expression of luciferase gene (*pMIR500A-Luc*, represented in the figure) and we studied its promoter activity in TERT-overexpression conditions by luciferase reporter assay (**C**). Then, we studied the effect of inhibiting *TERT* expression in HEK 293 cells by using a specific siRNA on the *MIR500A* promoter activity (**D**). Finally, we determined the promoter occupancy by the amplification of a ChIP assay in hTERT-SAOS 2 cells (**E**). The scheme represents the primers mapping to the MIR500 cluster. *TBE_cMyc* acts as a positive control and *intron_GAPDH* acts as a negative control. Each bar represents the mean ± SEM from triplicate samples. Graphs are representative (B, E) or the average (C, D) of three (N= 3) (B-D) or two (N= 2) (E) independent experiments. ns, not significant; *p<0.05; **p<0.01; ***p<0.001 according to Student’s *t-*test (B, C) and ANOVA followed by Dunnett’s multiple comparison test (D, E).

The luciferase reporter results prompted us to investigate the TERT occupancy of *MIR500A* promoter by ChIP experiments. TERT associated with the promoter region containing the region upstream *MIR500A* but failed to bind the upstream sequence of *MIR532*, which also contains several TBE (**Fig. 2E**). As expected, TERT also bind the TBE found upstream the oncogen *MYC*, as previously shown (3).

These results suggest that the whole MIR500 cluster could be regulated by TERT. RT-qPCR analysis of parental and TERT-expressing SAOS 2 revealed that TERT induces the expression of *MIR500A, MIR362, MIR500B* and *MIR502* (**Figs. 3C-3F**) but did not affect that of *MIR532* (**Fig. 3B**), which is located upstream and TERT is unable to bind its TBE.

**Figure 3:**
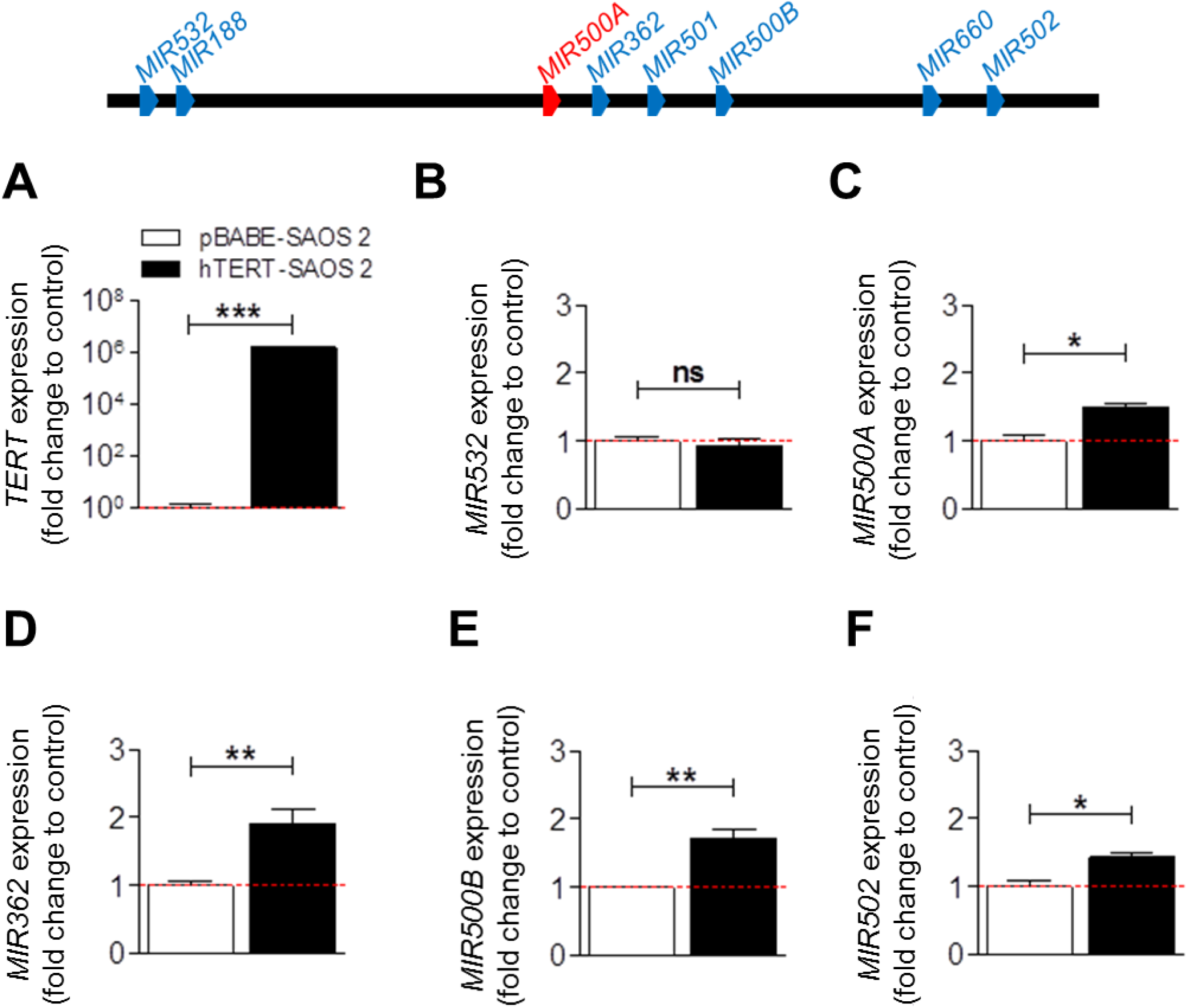
The MIR500 cluster is regulated by TERT. We studied the mRNA levels of five different *miRNA*s from the MIR500 cluster (**B**-**F**) under TERT-overexpression conditions (**A**) by real-time RT-qPCR. Each bar represents the mean ± SEM from triplicate samples and graphs are representative of three different experiments (N=3). ns, not significant; *p<0.05; **p<0.01; ***p<0.001 according to Student’s *t-*test.

### MIR500A mediates TERT-increased invasiveness of tumor cells

We next examined whether the different components of the cluster are also implicated in TERT-mediated tumor invasion. To achieve this goal, we overexpressed several miRNAs in parental and TERT-expressing SAOS 2 cells by transient transfection with the correspondent *pre-MIR* (**Fig. S4**) and studied the effect on the *in vivo* invasive capacity cells (**Fig. 4**). Surprisingly, *MIR500A* was the only one able to increase the *in vivo* invasion capacity of tumor cells, while the *MIR532*, which is not regulated by TERT, inhibited invasiveness of parental tumor cells (**Figs. 4B, 4D**). Collectively, these results show that *MIR500A* mediated *TERT*-induced tumor invasiveness.

**Figure 4:**
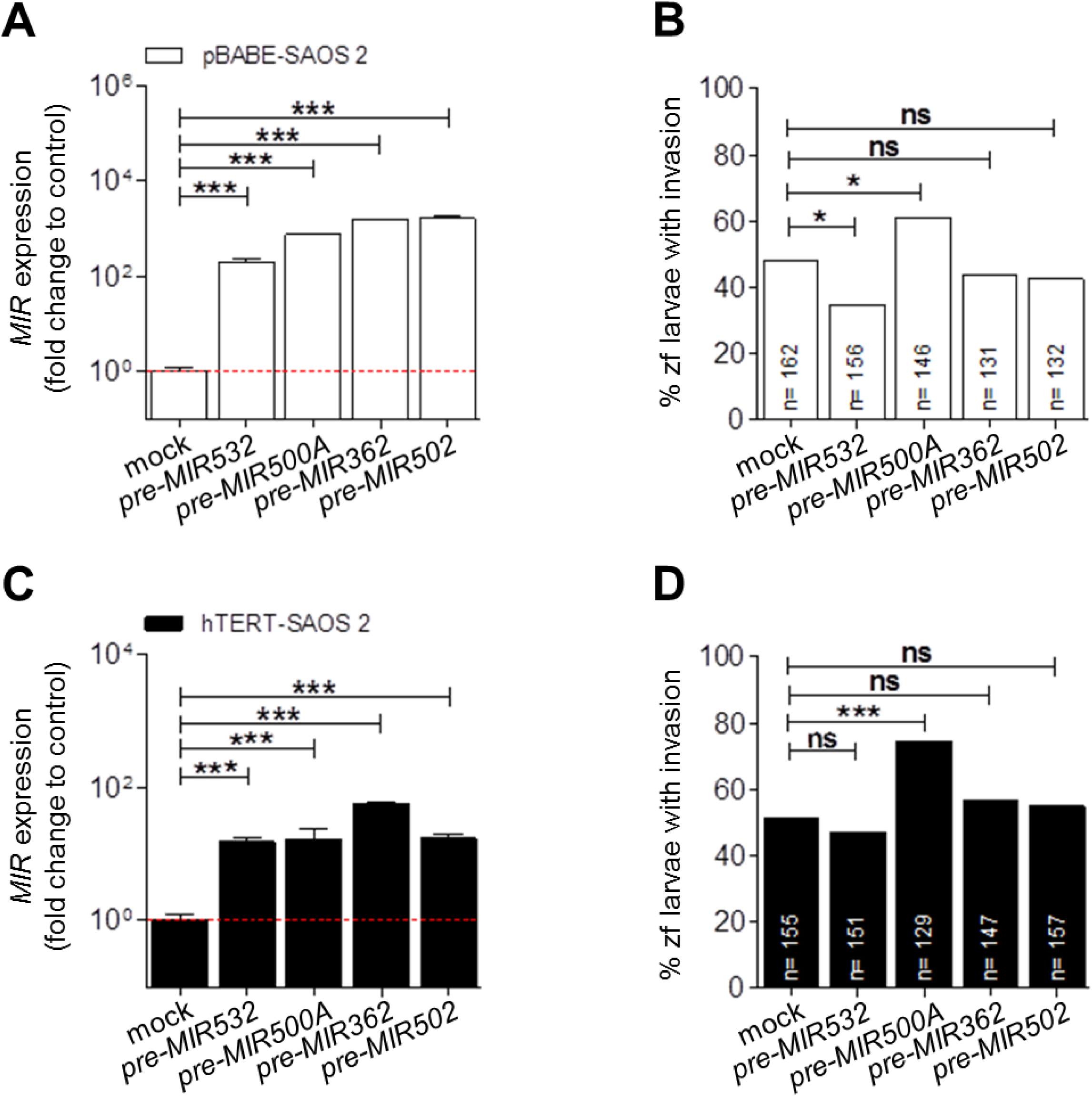
Only the *MIR500A* is able to increase the invasiveness. We studied the contribution of the different *miRNA*s from the MIR500 cluster to the *in vivo* invasion capacity of the pBABE-SAOS 2 (**A, B**) and the hTERT-SAOS 2 cells (**C**-**D**). In (A, C), each bar represents the mean ± SEM from triplicate samples. In (B, D), histograms represent the accumulative value of invasion percentage from a total larvae stated in the figure for each treatment. Graphs are representative (A, C) or the accumulative value (B, D) of three (N= 3) different experiments. ns, not significant; *p<0.05; ***p<0.001 according to ANOVA followed by Dunnett’s multiple comparison test (A, C) and Fisher’s exact test (B, D).

### The regulation of MIR500A by TERT does not depend on telomerase activity

To ascertain whether TERT requires its telomerase activity to regulate the expression of *MIR500A*, we used a dominant-negative mutant of TERT (DN-TERT), which has two point mutations in the A motif at the RT domain that cause it to lack the enzymatic activity (17). DN-TERT was expressed at the same levels than wild type TERT (**Fig. 5A**) and also increased *MIR500A* promoter activity (**Fig. 5B**), *MIR500A* transcript levels (**Fig. 5C**) and tumor invasiveness *in vivo* (**Fig. 5D**), in a similar level than TERT. To further confirm the novel non-canonical role of TERT, we used two different telomerase inhibitors: BIBR 1532, that binds and blocks TERT (18) and TAG 6, that binds and blocks *TERC* (19). Although both drugs were able to inhibit telomerase activity (**Fig. S5A**), only BIBR 1532 decreased both the expression of *MIR500A* (**Fig. 5E**) and the tumor invasiveness *in vivo* (**Fig. 5F**). As expected, and to discard any off-target effect, the treatment of parental SAOS 2 cells with these drugs did not affect either the expression of *MIR500A* (**Fig. S5B**) or the tumor invasiveness *in vivo* (**Fig. S5C**).

**Figure 5:**
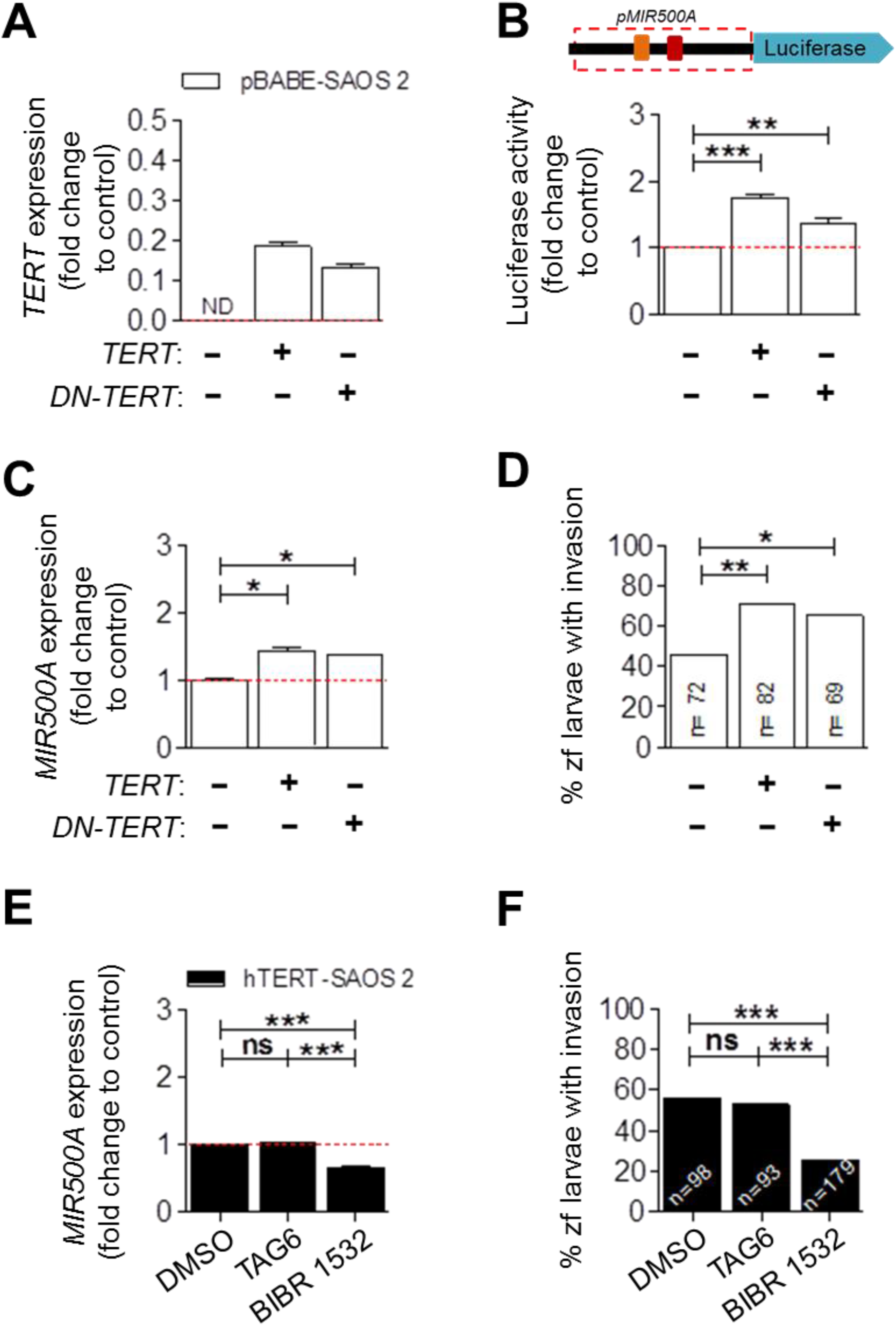
Telomerase activity is not involved in the *MIR500A* up-regulation by TERT. We co-transfected pBABE-SAOS 2 cells with *TERT* or *DN-TERT* (**A**-**D**) to determine whether telomerase activity is necessary or not for the *MIR500A* promoter activity (**B**), TERT-dependent *MIR500A* expression (**C**), for and for the *in vivo* invasive capacity (**D**). We also used two different chemical inhibitors (TAG-6 and BIBR 1532, which block *TERC* and TERT subunits, respectively) in hTERT-SAOS 2 cell line and we studied the drug effect on the levels of *MIR500A* (**E**) and on the *in vivo* invasive capacity (**F**). Each bar represents the mean ± SEM from triplicate samples and graphs are representative of three (N=3) independent experiments (A-C, E). Histograms represent the accumulated value of invasion percentage from a total larvae stated in the figure for each treatment and graphs are the average of two or three (N=2, =3) independent experiments (D, F), respectively. ns, not significant; *p<0.05; **p<0.01; ***p<0.001 according to ANOVA followed by Tukey’s multiple comparison test (B, C, E) and Fisher’s exact test (D, F).

### Hedgehog signaling pathway is regulated by MIR500A

The *Target Scan* software (*https://www.targetscan.org*) revealed that the 3’UTR of 3253 human genes contain putative target sites for *MIR500A*. By using the *MetaCore* software (*https://www.omictools.com/metacore-tool*), we classified the signaling pathways enriched in the predicted targets of *MIR500A* and found crucial role involved in cancer aggressiveness, such as Notch, WNT and Hedgehog signaling pathways (**Fig S6**). We focused in the latter, since *PTCH1, GLI3* and *CUL3* have all a putative target site for *MIR500A* (**Fig. 6A**). We confirmed by real-time RT-qPCR that *PTCH1, GLI3* and *CUL3* were all down-regulated in TERT-overexpression conditions (**Figs. 6B-6D, S7A**). In addition, *MIR500A* directly bound to *PTCH1* 3’UTR, as assayed by luciferase reporter experiments (**Figs. 6E, S7B**).

**Figure 6:**
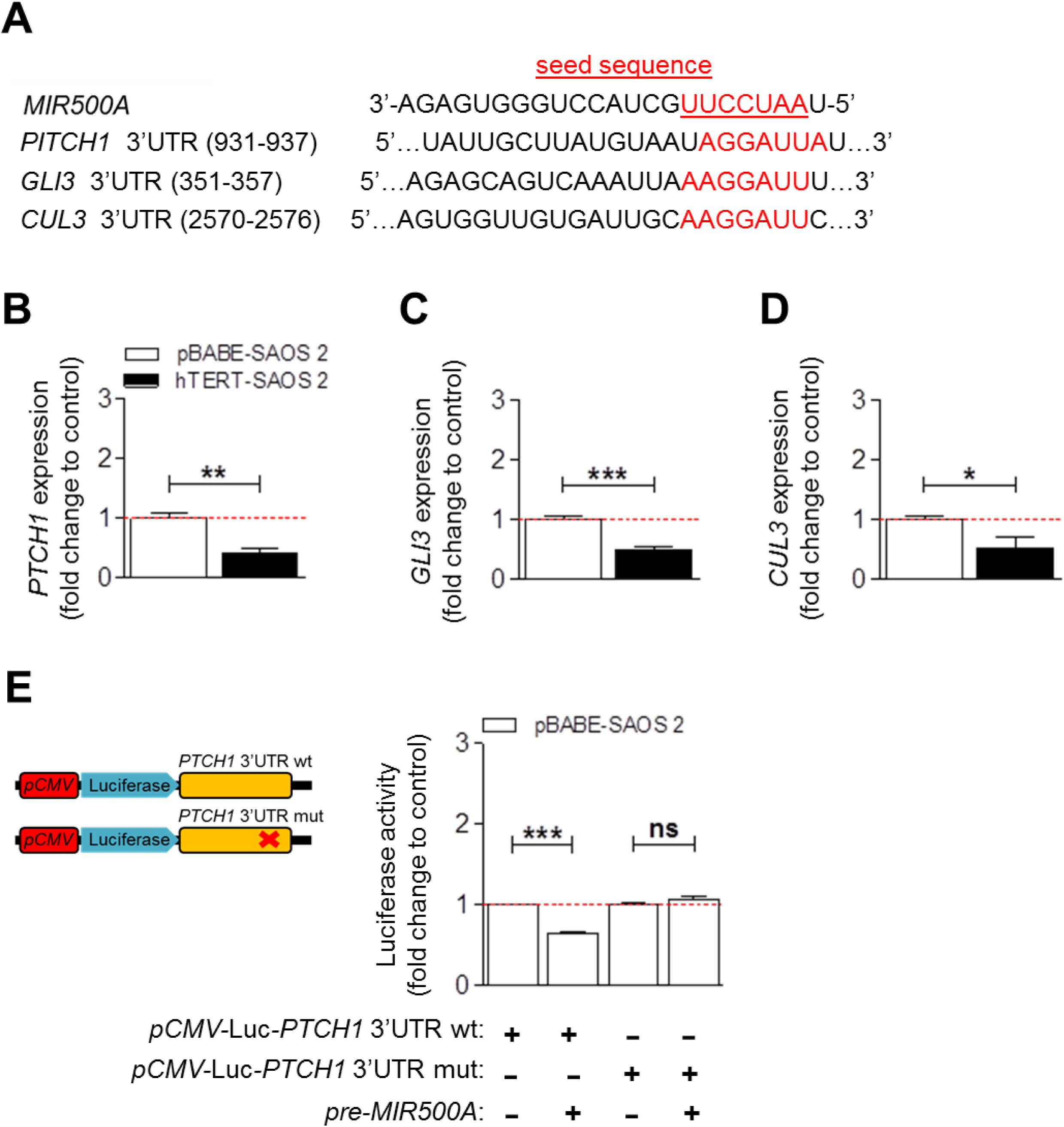
The Hedgehog signaling pathway is regulated by *MIR500A*. Alignment between the seed sequence of the *MIR500A* (underlined, in red) and the 3’UTR of *PTCH1, GLI3* and *CUL3*, from the Hedgehog signaling pathway (**A**). We studied the effect of *MIR500A* over the mRNA levels of *PTCH1* (**B**), GLI3 (**C**) and CUL3 (**D**) by real-time RT-qPCR. Finally, we validated the direct binding of *MIR500A* over the 3’UTR of *PTCH1* through luciferase experiments (**E**). Graphs are the mean of three (N=3) independent experiments (B-E). *p<0.05; ***p<0.001 according to Student’s *t* test (B, C, D) and ANOVA followed by Bonferroni’s multiple comparison test (E).

The above results prompted us to analyze if there is a correlation among *TERT, MIR500A* and *PTCH1* expression in cancer. We observed a significant positive correlation between the expression of TERT and *MIR500A* and a negative one between *PTCH1* and *MIR500A* in stomach adenocarcinoma and bladder urothelial carcinoma from *The Cancer Genome Atlas* (TCGA) cohort (*https://www.cancer.gov/tcga*) (**Figs. S7C, S7D**). These results point out to the relevance of inhibiting the extracurricular functions of telomerase in these specific cancer histotypes.

## Discussion

The identification and understanding of the non-canonical functions of TERT will provide new and important insights into the role of telomerase in cancer progression, helping in the development of specific strategies for the therapeutic manipulation of TERT in human cancer. Therefore, we decided to investigate if telomerase may regulate tumor invasion through the regulation of miRNA expression. To study exclusively the non-canonical functions of telomerase in this process, we generated a cell line model by stable transfection of the telomerase-negative cell line SAOS 2 with exogenous TERT. Taking advantage of the xenograft assay in zebrafish, we have validated our model by confirming that *TERT* overexpression increases the *in vivo* invasive capacity of hTERT-SAOS 2 compared with the parental cell line transfected with the empty plasmid (pBABE-SAOS 2).

As a starting point, to study the triad TERT-miRNA-invasiveness, we used a miRNA array approach to analyze miRNA expression changes in TERT-overexpression conditions. Surprisingly, the analysis showed a single statistically significant up-regulated miRNA, the oncomiR *MIR500A.* In our hands, the overexpression of *MIR500A* in both pBABE-SAOS 2 and hTERT-SAOS 2 increased tumor invasion in xenografted zebrafish larvae, indicating the implication of this miRNA in tumor invasion *per se*. Conversely, the inhibition of *MIR500A* decreased tumor invasiveness but, interestingly, only in the tumor cells that express TERT, pointing out to other players in the *in vivo* invasive capacity of SAOS 2 cells and demonstrating that TERT increases tumor invasiveness through *MIR500A*. The inhibition of *MIR500A* in another telomerase-positive cell line, HeLa 1211, resulted in a similar outcome, confirming that the identified mechanism operate in different tumor histotypes. The oncogenic activity of *MIR500A* is not surprising, since high serum level of *MIR500A* is a diagnostic biomarker of hepatocellular carcinoma (20), is associated with poor prognosis and overall survival in prostate cancer (21) and is also highly correlated with malignant progression and poor survival in gastric cancer (22).

The regulation of miRNAs is poorly understood, due in part to the difficulty in predicting promoters from short conserved sequences. We localized *MIR500A* in a cluster of 8 miRNAs: *MIR532, MIR118, MIR500A, MIR362, MIR501, MIR500B, MIR660* and *MIR502*. This cluster is into the short arm of the X chromosome (Xp11.23), in the intron 3 of the *CLCN5* gene. A few studies indicate that intronic miRNAs are not necessarily co-transcribed with their host gene, which suggests that they might have their own independent intronic promoters (23). Our results showed that the expression of *CLCN5* is not affected by the presence of TERT, which indicates that the regulation of *MIR500A* by TERT is not mediated through the regulation of its host gene promoter. As it has been reported that TERT directly interacts with TBE-containing promoters (3, 4), we decided to analyze the *MIR500A* upstream sequence. Notably, the sequence analysis revealed the presence of various TBE sequences upstream of *MIR500A* and *MIR532*. We found that TERT was able to increase the expression of a luciferase reporter driven by a 2 Kb fragment upstream of *MIR500A* while, conversely, telomerase inhibition by siRNA decreased luciferase activity. These results were further confirmed by ChIP assays, where TERT was found to bind directly to the upstream region of *MIR500A*, but not to the upstream region of *MIR532*, despite it also contains a TBE. In addition, TERT also regulated *MIR362, MIR500B* and *MIR502*, all downstream *MIR500A*. However, the expression level of *MIR532*, which is located upstream *MIR500A* is unaffected by TERT. Collectively, these results demonstrate that TERT behaves as a transcription factor that up-regulates the expression of *MIR500A* and all its downstream miRNAs.

According to the evolutionary model proposed by Chen and Rajewsky (24), a possible hypothesis to explain the TERT-dependent regulation of the MIR500 cluster, excluding the two miRNAs located upstream of *MIR500A*, is that although TERT originally could be interacting with a TSS (Transcriptional Start Site) region in a potential intronic promoter to regulate the expression of the whole cluster, *MIR500A* became the main effector of the cluster and developed its own TERT-regulated promoter, while *MIR532*, located upstream of this promoter, became a negative regulator of *TERT* as a compensatory mechanism to fine-tuning TERT levels. However, this negative feedback has to be confirmed with further experiments.

It has also been observed that genes involved in development have more transcription factor (TF)-binding sites and miRNA-binding sites on average, revealing that the genes with higher *cis*-regulation complexity are coordinately regulated by TFs at the transcriptional level and by miRNAs at the post-transcriptional level (25). Based on this observation, it is tempting to speculate that *TERT*, a crucial gene for life, regulates and is regulated by miRNAs of the same cluster. In line with this speculation, *MIR500A* and all downstream miRNAs of the cluster, act as oncomiRs and are related to different cancer types: *MIR500A* in hepatocellular carcinoma, gastric and breast cancer (26-28), *MIR362* in chronic myeloid leukemia (29), *MIR501* in gastric cancer (30), *MIR660* in breast cancer (31) and *MIR502* in colorectal and prostate cancer (32, 33). Conversely, the miRNAs located upstream *MIR500A* act as tumor suppressors: *MIR532* inhibits the expression of *TERT* in ovarian cancer, resulting in decreased cell proliferation and invasion capacity (34), and *MIR188* is down-regulated in oral squamous cell carcinoma (35). In addition to being transcribed together, the different miRNAs of a cluster usually have the same function. To explore this possibility, we studied the effect of different components of the *MIR500* cluster on the *in vivo* invasion capacity of SAOS 2 cells and, interestingly, only *MIR500A* was able to increase tumor invasion, while *MIR532* had the opposite effect. Altogether, these data highlight the relevance of TERT in the regulation of the MIR500 cluster and the relevance of this crosstalk in cancer progression.

To catalog as a non-canonical function the ability of TERT to regulate the MIR500 cluster and its relevance in cancer invasion, we studied whether the absence of telomerase activity affects this activity and the invasiveness of tumor cells *in vivo* by two different approaches: a genetic approach using DN-TERT, and a pharmacological inhibition of either TERT or *TERC* subunits. Surprisingly, neither DN-TERT nor chemical inhibition of *TERC* with TAG 6 affected any function apart of the telomerase activity *per se*, while BIBR 1532 reduced the expression of *MIR500A* and, consequently, decreased tumor invasiveness *in vivo*. These results support the hypothesis of an extracurricular role of TERT in transcriptional regulation of the MIR500 cluster through its direct binding to the genomic DNA, helping the cancer progression and metastasis. Furthermore, they also point to the importance of choosing the right strategy when using telomerase inhibitors against cancer, since it can be more important to inhibit the extracurricular role of TERT by physically preventing its binding to DNA than inhibiting its enzymatic activity.

*Target Scan* database prediction and the *MetaCore* software functional annotations of predicted targets revealed crucial signaling pathways downstream the TERT/MIR500 cluster axis, such as Wnt/β-catenin, NF-κB and Hedgehog, among other. We focus our attention on the Hedgehog (Hh) signaling pathway, since it is well established that its aberrant activation leads to enhanced proliferation and invasion of tumor cells and we found that TERT-induced *MIR500A* mediated the down-regulation of *PTCH1, GLI3* and *CUL3*, and *MIR500A* directly targets the 3’UTR of *PTCH1*, promoting tumor invasiveness. The function of the receptor of Hh signaling pathway PTCH1 as a tumor suppressor is not surprising and has already been shown in other studies (36, 37).

By using the data generated by the TCGA Research Network, we have found a strong positive correlation between *TERT* and *MIR500A*, while *MIR500A* expression was found to be negatively correlated with that of *PTCH1* in stomach adenocarcinoma and bladder urothelial carcinoma. A high-stage of gastric cancer and negative PTCH1 staining have been identified as unfavorable risk factor for overall survival (438) and it has been described an important role of Hh signaling in bladder cancer growth and tumorigenicity (39). In addition, Shh signaling crosstalks with other signaling pathways during development and cancer progression, such as Notch, Wnt, and TGF-β signaling pathways (40), which are also regulated by *MIR500A* (26) and TERT (6). Therefore, our results support that the *MIR500A* is also a good therapeutic target to fight cancer.

In summary, we have demonstrated for the first time that TERT is able to regulate the expression of specific microRNAs through its direct binding to TBE regions at their promoter sequence. In particular, TERT-mediated up-regulation of *MIR500A* results in a post-transcriptionally repressesion of *PTCH1*, which triggers a ligand-independent aberrant Hh signaling activation that significantly increases tumor cell invasiveness in a zebrafish xenograft model (**Fig. 7**). This is a novel non-canonical telomerase function, since is independent of telomerase activity, paving the way in the development of new therapeutic strategies to fight cancer through the inhibition of extracurricular activities of TERT.

**Figure 7:**
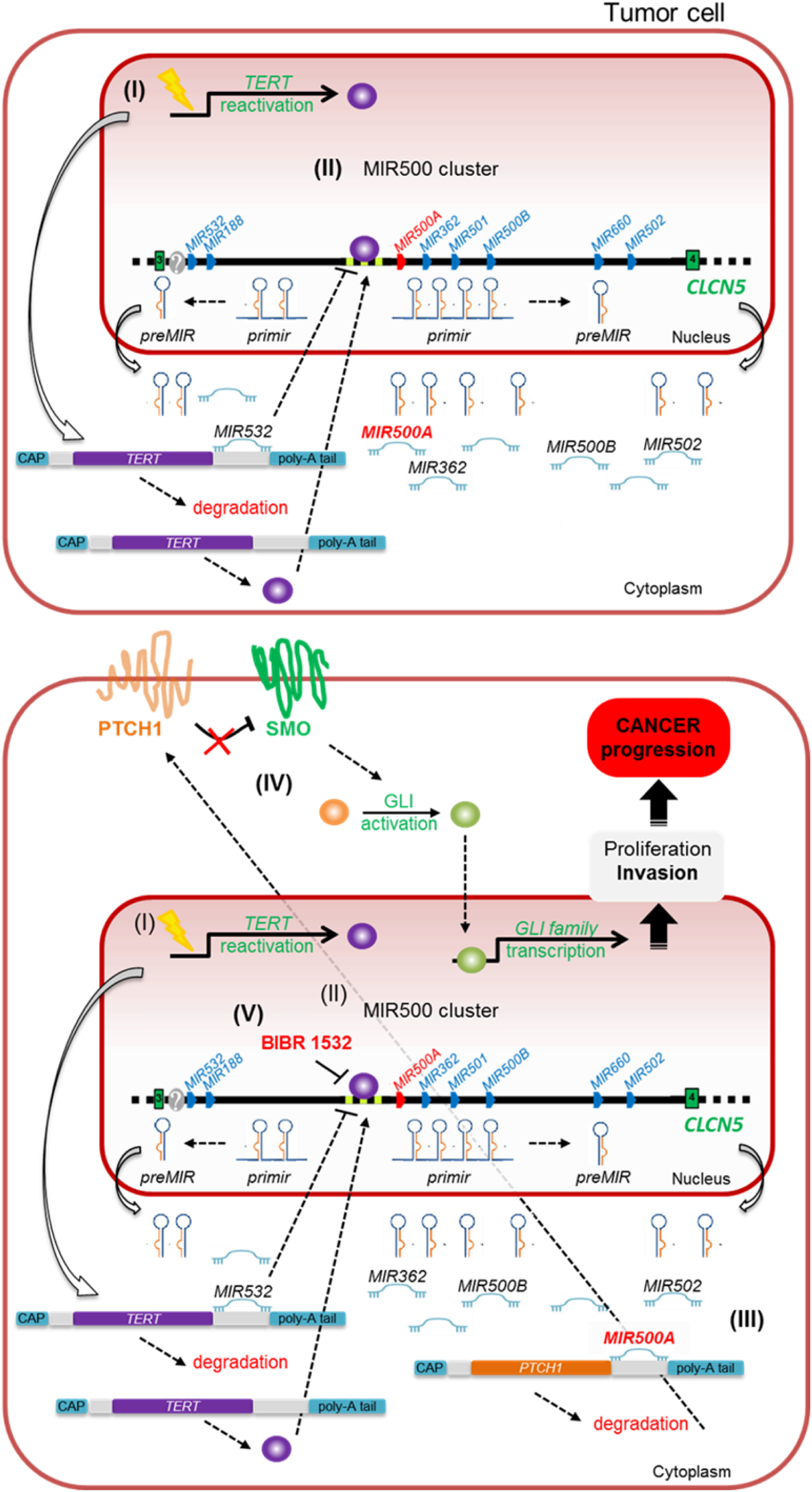
Extracurricular mechanism of TERT in invasion and tumor progression through the regulation of *MIR500A*. (**I**) Telomerase expression is reactivated in most tumors and (**II**) TERT binds directly to the TBE sequences located at the promoter region of *MIR500A*, resulting in the up-regulation of *MIR500A* and also the miRNAs located downstream it. As a compensatory mechanism, this regulation is fine-tuned by the *MIR532*, which acts as a negative regulator by repressing the *hTERT* mRNA. (**III**) The oncomiR *MIR500A* represses post-transcriptionally the mRNA of the tumor suppressor *PTCH1*, triggering a ligand-independent aberrant Hedgehog signaling activation (**IV**) that contributes significantly to increase the invasiveness of tumor cells. (**V**) The chemical inhibition of TERT with BIBR 1532 could be a new strategy to fight cancer.

## Methods

### Animals

Zebrafish (Danio rerio H., Cypriniformes, Cyprinidae) were obtained from the Zebrafish International Resource Center and mated, staged, raised and processed using standard procedures. Details of husbandry and environmental conditions are available on protocols.io (DOI:dx.doi.org/10.17504/protocols.io.mrjc54n).

The experiments performed comply with the Guidelines of the European Union Council (86/609/EU). Experiments and procedures were performed as approved by the Bioethical Committee of the University Hospital “Virgen de la Arrixaca” (HCUVA, Spain).

### Cell culture

Human embryonic kidney 293 (HEK 293), cervical cancer (HeLa 1211) and sarcoma osteogenic (SAOS 2) cell lines were purchased from the ATCC (#CRL-1573.3, #CCL-2 and #HTB-85, respectively). All cell lines were maintained in DMEM (Sigma, #D-5796) supplemented with 10% FBS (Biowest, #S1810-500), and were cultured at 37 °C with 5% CO_2_.

pBABE-SAOS 2 and hTERT-SAOS 2 stable cell lines were obtained upon transfection of the SAOS 2 cell line with the plasmids pBABE-puro or pBABE-puro-hTERT from Addgene (#1764, #1771, respectively), and Lipofectamine 2000 (Invitrogen, #11668-027) following manufacturer’s instructions. Then, several stable clones were selected with puromycin.

### Zebrafish xenograft assay

Cells were trypsinized, washed and stained with the vital cell tracker red fluorescent CM-Dil (4 ng/ul final concentration, Invitrogen, #C7001), following manufacturer’s instructions. Zebrafish larvae, previously treated with PTU (N-phenylthiourea, Sigma-Aldrich, #222909) to inhibit the skin pigmentation, were dechorionated and anesthetized with tricaine (Sigma, #A5040). Then, 100-150 labelled cells in 4 nl were injected into the yolk sac of 2 dpf zebrafish larvae using a manual injector (Narishige IM-300, East Meadow (Long Island), NY, USA). After injection, embryos were incubated for 2 h at 31 °C and checked for cell presence at 2 hours post-xenograft (hpx). Fish with fluorescent cells outside the implantation area at 2 hpx were excluded for further analysis. All other fish were incubated at 35 °C for 48 h and analyzed with a SteReo Lumar.V12 stereomicroscope with an AxioCam MR5 camera (Carl Zeiss, Thornwood, NY, USA). Evaluation criteria for invasion were that at least three cells had to be identified outside the yolk.

### miRNA microarray

RNA from 18 nucleotides (nt) upwards was isolated from two different clones of pBABE-SAOS 2 and hTERT-SAOS 2 stable cell lines by using miRNeasy Mini Kit (Qiagen, #79306), following manufacturer’s instructions. A total of 0.5 µg of RNA from each sample were sent to the CNIO Genomic Facility. For quality control, all samples were analyzed on a Nanodrop instrument (Bioanalyzer 2100, Agilent) by the Facility. There, they were labeled and hybridized on the Human miRNA 8×15K, 1 color array (Agilent, #G4470C). These arrays contained probes for 2689 microRNAs. Data analysis was performed and we obtained a gene list according to a *P*-value. Only one gene had a *P*-value<0.05 (*MIR500A*).

### Gene and microRNA expression analysis

RNA from 18 nucleotides (nt) upwards was extracted from 10^6^ cells homogenized in QIAzol Lysis Reagent (Qiagen, #79306) and using the miRNeasy Mini kit (Qiagen, # 217004), following manufacturer’s instructions. cDNA was generated by the miScript II RT kit (Qiagen, #218161), following the manufacturer’s instructions, and treated with DNase I, amplification grade (1 U/µg RNA, Qiagen, #79254). Real-time qPCR was performed with a MyiQ™ instrument (BIO-RAD), using miScriptSYBR Green PCR kit (Perfect Real Time) (Qiagen, #218161). Reaction mixtures were incubated for 10 min at 95 °C, followed by 40 cycles of 15 s at 95 °C, 1 min at 60 °C, and finally 15 s at 95 °C, 1 min at 60 °C, and 15 s at 95 °C. For each sample, microRNA or gene expression was normalized to *U6* snRNA or *GAPDH* content in each sample, respectively, using the comparative *Ct* method (2^-ΔΔ*Ct*^). The primers used are shown in supplementary **Table S1**. In all cases, each PCR was performed with triplicate samples and repeated, at least, with two independent samples.

### Overexpression experiments

*TERT* (Addgene plasmid #1771) and a dominant-negative mutant (*DN-TERT*) (Addgene plasmid #1775), and different members of the MIR500 cluster (*pre-MIR532, -500A, -362* and *-502*, from Ambion, #PM11553, #PM12793, #PM10870, #PM10480, respectively) were overexpressed in pBABE-SAOS 2 and hTERT-SAOS 2 upon transfection with Lipofectamine 2000 following manufacturer’s instructions. 48 hours after transfection, cells were trypsinized and divided to functional assays and to measure the expression level by real time RT-PCR analysis.

### Silencing experiments

To inhibit the *MIR500A*, a specific Peptide Nucleic Acid (PNA) miRNA inhibitor was used (Panagene, #PI-1487-FAM). After a 10 min incubation at 70 °C in a water bath or heating block of the PNA miRNA inhibitor, cells were transfected with a final concentration of 500-2,000 nM by using Lipofectamine 2000, following manufacturer’s instructions.

For telomerase silencing, HEK 293 cells were transfected with a ready-to-use siRNA for human TERT (*TERT siRNA (h)*, Santa Cruz Biotechnology, #sc-36641) at a final concentration of 20 mM by using Lipofectamine 2000, according to manufacturer’s instructions. 48 h after transfection, cells were trypsinized and divided to functional assays and to measure the knock-down efficiency by real time RT-PCR analysis.

### Analysis of MIR500A promoter activity

A 2 Kb genomic DNA sequence upstream of *MIR500A* +1 position was amplified using the primers: forward 5’CAGTGTTGTGGTTTTGGTCCAGGCG3’ and reverse 5’CCGGACACCGAGCACCGGCGAGCCGCC3’. The DNA fragment was cloned in the *SmaI* site of the pGL3basic vector (Promega, #E1761) driving the expression of firefly luciferase reporter gene (*pMIR500A-*Luc). Cells were transfected with a mix containing 100 ng/cells the firefly luciferase construct and 50 ng/µg of *Renilla* luciferase control plasmid by using Lipofectamine 2000, according to manufacturer’s instructions. After 48 h, cell extracts were obtained and assayed for luciferase activity by using the Dual-Luciferase assay kit (Promega, #E1910), as specified by the manufacturer, in an Optocomp I luminometer (MGM Instruments).

### Chromatin immunoprecipitation assay, ChIP

Both pBabe-SAOS 2 and hTERT-SAOS 2 (10^7^ cells) were cross-linked with 1% paraformaldehyde (Sigma Aldrich, # P6148) in culture medium for 10 min at room temperature. Then, aldehydes were quenched with PBS containing 200 mM glycine (Sigma Aldrich, #M6635) for 5 min followed by a PBS wash. The cells were centrifuged at 200 xg for 10 minutes at 4 °C to pellet. Then, they were resuspended in Lysis Buffer containing Protease Inhibitors (Sigma-Aldrich, #P2714) and the lysate was sonicated using a sonication system Bioruptor Plus (Diagenode) for 30 cycles of 30 seconds ON, 30 seconds OFF. The sonicated lysate was centrifuged at 20,000 xg for 10 min at 4 °C, and the supernatant was transferred to new tubes. ChIP dilution buffer (100 uL) was added to the supernatant, and 10 uL of the supernatant was set aside for input. Binding the chromatin to the Antibody-Dynabeads complexes, reverse the formaldehyde crosslinking of the chromatin and purifying the DNA were performed as described in the protocol MAGnif Chromatin Immunoprecipitation System (Invitrogen, #49024).

The results of the ChIP were analyzed by real-time qPCR, with a qPCR ABI PRISM 7500 instrument (Applied Biosystems), using the commercial kit Power SYBR Green PCR Master Mix (Applied Biosystems, #4309155). Reaction mixtures were incubated for 15 min at 95 °C, followed by 40 cycles of 15 s at 94 °C, 30 s at 55 °C, and finally 30 s at 70 °C, 1 min at 95 °C, 1 min at 60 °C. The primers used are shown in supplementary **Table S2**.

### Quantitative telomerase activity assay, Q-TRAP

To quantitatively measure the telomerase activity, total proteins were extracted from cells using ice-cold CHAPS lysis buffer (Sigma-Aldrich, #S7705) and real-time Q-TRAP performed with 0.1 μg protein extracts. A negative control of each sample confirmed the specificity of the assay (data not shown in figure).Control samples were obtained by treating the cell extracts with 1 μg RNase (ThermoFisher, #EN0531F) at 37 °C for 20 min. For making the standard curve, a 1:10 dilution series of telomerase-positive sample (HeLa cells) was used. After qPCR amplification, real time data were collected and converted into Relative Telomerase Activity (RTA) units performing the calculation: RTA of sample = 10^(Ct sample-Yint)/slope^. The standard curve obtained was: y= 23.802–3.2295x.

### Chemical inhibition of telomerase activity

BIBR 1532 (Santa Cruz Biotechnology, #sc-203843) and TAG-6 (Calbiochem, #581004) were added to 20 µM and 2.5 µM final concentration in cell culture, respectively. As a control, DMSO was adjusted to the same concentration. Cells were incubated with the compounds during 15 h, then trypsinized and divided for xenografting, checking telomerase activity by Q-TRAP and studying gene expression by real time RT-PCR.

### MIR500A target prediction and validation

We have used the *Target Scan* database (*http://www.targetscan.org*) to predict the potential targets. Then, we validated the chosen targets by real-time RT-qPCR and luciferase experiments. To validate the specific binding of *MIR500A* to the *PTCH1* 3’UTR, a 1.2 Kb genomic DNA 3’UTR sequence of *PTCH1* was amplified using the primers: forward 5’AAGGTCTAGAGCAAAGAGGCCAAAGATTGGA3’ and reverse 5’TCTAGAAAGCCTCAACCAGC3’. We also amplified the same region but lacking the *MIR500A* binding site using the primers: forward 5’AATATTGCTTATGTAA TATTATTTTGTAAAGG3’ and reverse 5’CCTTTACAAAATAATATTACATAAGCAATATT3’. The 3’UTR fragments were cloned in the *XbaI* site of the pRL-CMV vector (Promega, #E2261) (*pCMV-*Luc-*PTCH1* 3’UTR wt/mut). Cells were transfected and luciferase experiments performed as explained before.

### Statistical analysis

All data are expressed as the mean ± standard error of mean (SEM). Values of p<0.05 were considered statistically significant. Statistical analyses were analyzed using analysis of variance (ANOVA) followed by different post-hoc comparison tests. The differences between two samples were analyzed by the Student’s *t-*test. The percentage of zebrafish larvae with invasion was analyzed by chi square (Fisher’s exact test). All analyses were performed with GraphPad Prism 5.

## Acknowledgements

We strongly thank María C. López-Maya for their excellent technical assistance. The plasmid pBABE-puro was a gift from Hartmut Land, Jay Morgenstern and Bob Weinberg and the plasmids pBABE-puro-hTERT and pBABE-puro-DN-hTERT were a gift from Bob Weinberg.

## Author Contributions

MLC conceived the study; MBG, BRN, DGM, JGC, FAP, VM and MLC designed research; MGB, EMB, DGM, JGC, BRN, EBA, performed research; MGB, EMB, DGM, JGC, BRN, EBA, VM, FAP and MLC analyzed data; and FAP and MLC wrote the manuscript with minor contribution from other authors.

## Data and materials availability

All data needed to evaluate the conclusions in the paper are present in the paper and/or the Supplementary Materials. Correspondence and request for materials should be addressed to MLC or FAP.

